# Exendin-4 improves neuron protection and functional recovery in experimental spinal cord injury in rats through regulating PCBP2 expression

**DOI:** 10.1101/2020.11.09.373993

**Authors:** Huaichao Luo, Qingwei Wang, Lei Wang

**Affiliations:** Department of Clinical Laboratory, Sichuan Cancer Hospital & Institute, Sichuan Cancer Center, School of Medicine, University of Electronic Science and Technology of China, Chengdu, 610041, China; The Sichuan Provincial Key Laboratory for Human Disease Gene Study, Hospital of the University of Electronic Science and Technology and Sichuan Provincial People’s Hospital, Chengdu, 610072, China; Department of Medical Ultrasound, Sichuan Academy of Medical Sciences & Sichuan Provincial People’s Hospital, Chinese Academy of Sciences Sichuan Translational Medicine Research Hospital; Chengdu, 610072, China

**Author notes:** Corresponding Author: Lei Wang, Department of Medical Ultrasound, Sichuan Academy of Medical Sciences & Sichuan Provincial People’s Hospital, Chinese Academy of Sciences Sichuan Translational Medicine Research Hospital, No.32, West 2nd section, Yihuan road, Chengdu 610072, Sichuan, China.

**Keywords:** spinal cord injury, exendin-4, PCBP2, neuroprotective effect, cell apoptosis, inflammation

## Abstract

**Aims:** In the present research, we assessed the therapeutic effects of Exendin-4 (Ex-4) on rat models with spinal cord injury (SCI).

**Materials and methods:** 36 male Sprague–Dawley rats were randomly allocated into three groups, including sham operation group, SCI group and SCI+Ex-4 group (Ex-4 treatment (10 µg/rat) after SCI, i.p.). In the SCI group, a laminectomy was performed at the T10 vertebrae, followed by weight-drop contusion of the spinal cord. In the sham group, a laminectomy was carried out without SCI contusion.

**Key findings:** Our results showed that Basso-Beattie-Bresnahan scale scores were significantly decreased after SCI, and were obviously improved in SCI rats with Ex-4 administration. Additionally, the water content of spinal cord in SCI group was dramatically increased than that in sham group, and after Ex-4 treatment, degree of edema of spinal cord was remarkably reduced. And also, concentration levels of inflammatory cytokines (IL-1α, IL-1β, IL-6 and TNF-α) in the spinal cord were significantly elevated after SCI, and were remarkably reduced in SCI rats with Ex-4 administration. Subsequently, cell apoptosis rate in the injured spinal cord was significantly increased, and after Ex-4 treatment, cell apoptosis rate was remarkably decreased. We also revealed that levels of PCBP2 mRNA and protein were significantly up-regulated after SCI, and were dramatically dropped in SCI rats with Ex-4 administration.

**Significance:** Take altogether, our findings disclosed that Ex-4 plays a role in promoting neurological function recovery and inhibiting neuronal apoptosis through effecting PCBP2 expression in SCI rat models.

## Introduction

The pathophysiological changes of spinal cord injury (SCI), a highly devastating pathology that seriously harms human health worldwide (1), is categorized into two temporally-related mechanisms: the primary injury and the secondary injury. The primary injury refers to a short-time impairment caused by the initial mechanical trauma, which leads to irreversible damage of neurons. The secondary injury, induced by multiple biological events including oxidative stress, immune dysfunction and neuronal apoptosis (2-5), often causes a great number of neurological, behavioral, emotional, and cognitive deficits. Despite great achievements in the therapeutic approaches, the prognosis of SCI patients still remains poor due to severe neurological deficits. Accordingly, it is necessary for us to explore the main mechanisms of neuronal apoptosis in SCI and identify a novel anti-apoptotic drug for SCI treatment.

Oxidative stress is one of the important factors that cause the damage of SCI neurons. Oxidative stress response can cause the levels of reactive oxygen species and inflammatory factors increase, thereby inducing neuronal damage and apoptosis (6) (7). Nrf-2 / OH-1 is an antioxidant pathway that plays an important role in the body’s antioxidant response, inhibition of the Nrf-2 / OH-1 pathway will aggravate SCI, and activate the Nrf-2 / OH-1 pathway play a protective effect on neurons (8, 9). It is showed that Nrf-2 / OH-1 pathway can be activated by Exendin-4 (Ex-4) (10). Ex-4, a 39-amino acid peptide originally extracted from helodermasuspectum venom (11), is extensively considered to be an effective drug for treating diabetes in the past decades (12-14). In addition, more and more investigations showed that Ex-4 might serve a neuro-protective role in several neurodegenerative diseases, including amyotrophic lateral sclerosis (15), Huntington’s disease (16), Parkinson disease (17) and Alzheimer’s disease (18). However, recent study demonstrated that Ex-4 could prevent against SCI-induced mitochondrial apoptotic pathway (19). However, the physiological or pathological functions of Ex-4 on SCI remained to be further clarified.

PCBP2, a member of the poly(C)-binding protein (PCBP) family, exerts a crucial role in posttranscriptional and translational regulation through interacting with single-stranded poly(C) motifs in target mRNAs (20). Aberrant PCBP2 expression was closely associated with a wide variety of human diseases, such as hepatic insulin resistance (21), and cardiomyocyte hypertrophy (22). Roychoudhury et al. (23) showed that PCBP2 was down-expressed in oral cancer cells, and up-regulation of PCBP2 could contribute to a significant increase in the number of apoptotic cells. Moreover, previous evidence indicated that PCBP2 might play important roles in neuronal apoptosis and astrocyte proliferation after SCI (24).

In the present article, we aimed to explore whether Ex-4 could exert a therapeutic effect in SCI through regulating PCBP2 expression and Nrf-2 / OH-1 pathway. Our research might provide a potential therapeutic strategy for patients with SCI.

## Materials and methods

### Animals and groups

A total of 36 adult male Sprague–Dawley (SD) rats weighing from 250 to 300 g were purchased from Animal Center of Chinese Academy of Sciences, Shanghai, China. Animals were housed in individual cages under standard laboratory conditions with a 12-hour light/dark cycle and were given *ad libitum* access to standard diet and water. The rats were randomly divided into three groups (n=12 per group), including (1) sham operation group, (2) SCI group and (3) SCI+Ex-4 group. The rats in sham operation group were only received opening laminectomy operation and administrated with normal saline, the rats in SCI group were received the process of SCI and administrated with normal saline and the rats in SCI+Ex-4 group were administrated intraperitoneally (i.p.) with Ex-4 (10 µg/rat) (Sigma-Aldrich, St. Louis, MO, USA) in normal saline after SCI and dosing interval was 24h for three days. After 7 days, all rats were sacrificed through cervical dislocation to collect samples for further experiments. All procedure and handling techniques were approved by the Ethics Committee of Sichuan Provincial People’s Hospital and were in strict accordance with the recommendations in the National Institutes of Health Guide for the Care and Use of Laboratory Animals.

### Establishing SCI model

The rat models of SCI in this study were established based on the method by Allen as described previously (25). All rats in this study were anesthetized with intraperitoneal injection of 10% chloral hydrate (0.3 ml/kg, i.p.) and placed in a stereotactic frame. The skin and muscle were incised and a laminectomy was carried out to explore the thoracic vertebra level 10 (T10) of rats. Then a 10-g weight impactor was vertically dropped from a 20-mm height onto the exposed cord. After the operation above, the incision were closed with suture. In addition, the rats in sham operation group were only received the laminectomy without SCI. All animals were then placed on a warm pad until they recovered from anesthesia. Postoperative care included antibiotic (crystalline penicillin 80000 units diluted in 5 ml NS) injections twice a day for 7 days and manual bladder expression twice a day until spontaneous voiding occurred or till the end of the study.

### Assessment of behavior

The motor functions of rats, including limb function, the degree of coordinated stepping, stepping ability and trunk stability, were evaluated through the Basso, Beattie and Bresnahan (BBB) rating scale (26). The BBB scores range from 0 points, which indicates no observable hind limb movement, to 21 points, which indicates normal hind movement. Each part was carried out for 10 minutes through two individuals who were blind to treatment at pre-injury and 1, 3 and 7 days after surgery.

### Wet/dry weight ratio of spinal cord

3-mm spinal cord sections from the center of the injury sites of three groups were obtained and weighted for wet weight (WW). Then, all samples were dried under 110°C for 24 h and weighted for the dry weight (DW). [(WW − DW) × 100]/WW was considered to be the water content of spinal cord.

### Protein extraction and ELISA analysis

Following the instructions, the expression of inflammatory cytokines in spinal cord tissue of rats, including IL-1α, IL-1β, IL-6 and TNF-α, were detected through performing ELIS Aassay (Biotechnology Co., Ltd. Shanghai enzyme research, Shanghai, China). With the instruction, total proteins were extracted from spinal cord tissues of all groups by using protein extraction kit (Beyotime Biotechnology, Shanghai, China), and the protein concentration was detected using the Bicinchoninic Acid Protein Assay. Standard (50 μl) was set up and samples were added to the ELISA plate (40 μl dilution+10 μl sample). OD values detected by MTP-800 Microplate reader (Corona Electric, Tokyo, Japan). The cytokine contents were presented as pg/mg protein.

### RNA extraction and RT-qPCR analysis

According to the instruction, total RNA was extracted from spinal cord samples through using Trizol Reagents (Ke Min Biological Technology Co., Ltd., Shanghai, China). For analysis of PCBP2 mRNA expression, total RNA was reverse transcribed to cDNA using PrimeScript RT-PCR Kit (Takara, Dalian, China). All primers used in this study were recorded in **Table 1**. qPCR analysis was performed using SYBR Premix Ex Taq (Takara) on a 7900 Fast Real-Time PCR system (Applied Biosystems, Foster City, CA).The relative expression of PCBP2 mRNA was normalized to glyceraldehyde-3-phosphate dehydrogenase (GAPDH) mRNA and calculated using the 2^−ΔΔCt^ method (ΔCt = Ct^target gene^ – Ct^internal control^) (27).

**Table 1.**
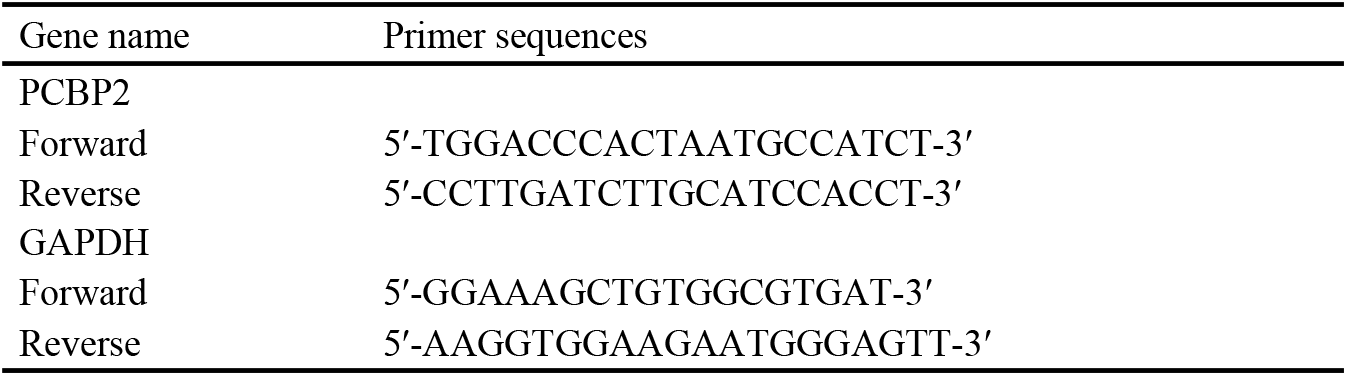
The sequences of primers.

### Western blot assay

Proteins were separated by 10% SDS-polyacrylamide gel electrophoresis, and transferred onto a nitrocellulose membrane (Bio-Rad, Shanghai, China). After blocking for 1 h, the membrane was incubated with primary antibodies (anti-PCBP2, 1:5000; Abcam, San Francisco, USA; anti-Nrf-2, 1:500; anti-OH-1, 1:500) for 4 h, and then incubated with goat-anti rabbit HRP-conjugated secondary antibody for another 2 hours. Immunoreactive protein bands were visualized using an ECL detection system (Amersham, Little Chalfont, UK) and gray values were analyzed by the Luminescent Image Analyzer (LAS-4000 mini, Fujifilm, Uppsala, Sweden). Relative optical density was calculated relative to GAPDH.

### TUNEL assay

Rats were anesthetized by intraperitoneal injection of about chloral hydrate (4%) and the rats were sacrificed. Spinal cord tissue was collected and embedded in paraffin. The sample was cut into 4 μm-thick slices, dewaxed with xylene, and incubated with gradient of ethanol. The samples were incubated with the proteinase K working solution for 30 min at 37 °C. The reagents were added in the dark according to the instructions of the In Situ Cell Death Detection Kit (Roche, US). Images were acquired under a fluorescence microscope.

### Statistical analysis

The data were presented as mean ± standard deviation (SD). All the data were analyzed and calculated by using SPSS 19.0 software (SPSS, Chicago, IL, USA) and GraphPad Prism 6 (GraphPad Software, La Jolla, CA, USA). One-way ANOVA and t-tests were used for the statistical analysis. The statistical tests were two-sided; a *P* value <0.05 was considered to indicate a statistically significant result.

## Results

### Exendin-4 protects hindlimb locomotion function after SCI

As shown in **Figure 1**, prior to model establishment, there was no remarkable difference in BBB locomotor rating scale scores in all groups. After SCI, the BBB scores in SCI group were significantly dropped in comparison to those in sham operation group, and the difference had statistically significant (all *P*<0.001). Moreover, after Ex-4 treatment, the BBB scores were obviously increased at 3 and 7 days after SCI in SCI+Ex-4 group compared with those in SCI group, and the difference had statistically significant (all *P*<0.05).

**Figure 1.**
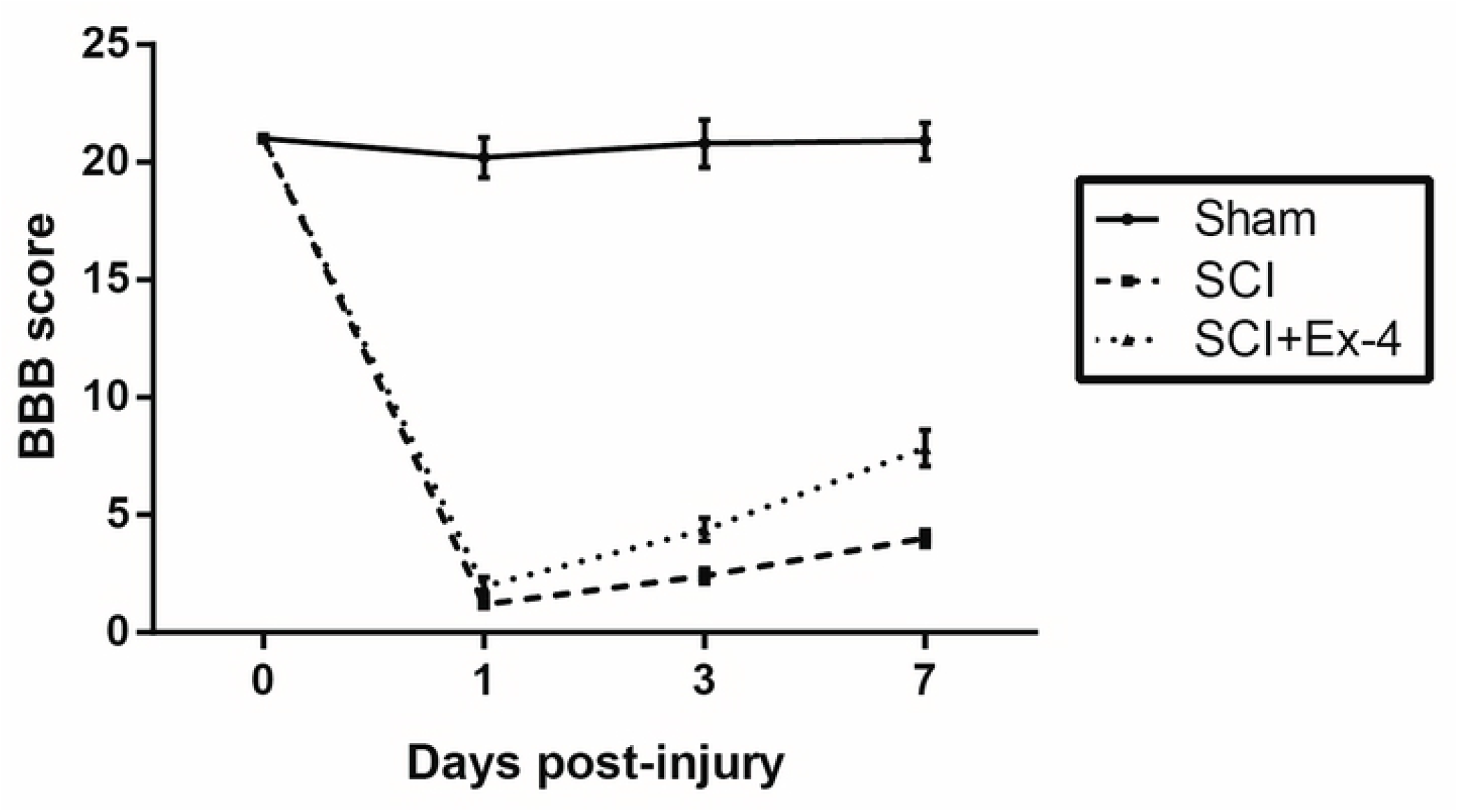
BBB locomotor rating scale in the three groups of rats, including a sham group, a SCI group and a SCI+Ex-4 group (n=12 per group). The BBB scores were obviously reduced in SCI group and were up-regulated after Ex-4 treatment.

### Exendin-4 alleviates edema of the injured spinal cord after SCI

To investigate the degree of edema of spinal cord in three groups, wet/dry weight ratio test was performed to calculate the water content of spinal cord in this study. As shown in **Figure 2**, the water content of spinal cord in SCI group was significantly elevated than that in sham group (*P*<0.01). After Ex-4 treatment, degree of edema of spinal cord was remarkably reduced in SCI+Ex-4 group in comparison with that in SCI group, and the difference also had statistically significant (*P*<0.05).

**Figure 2.**
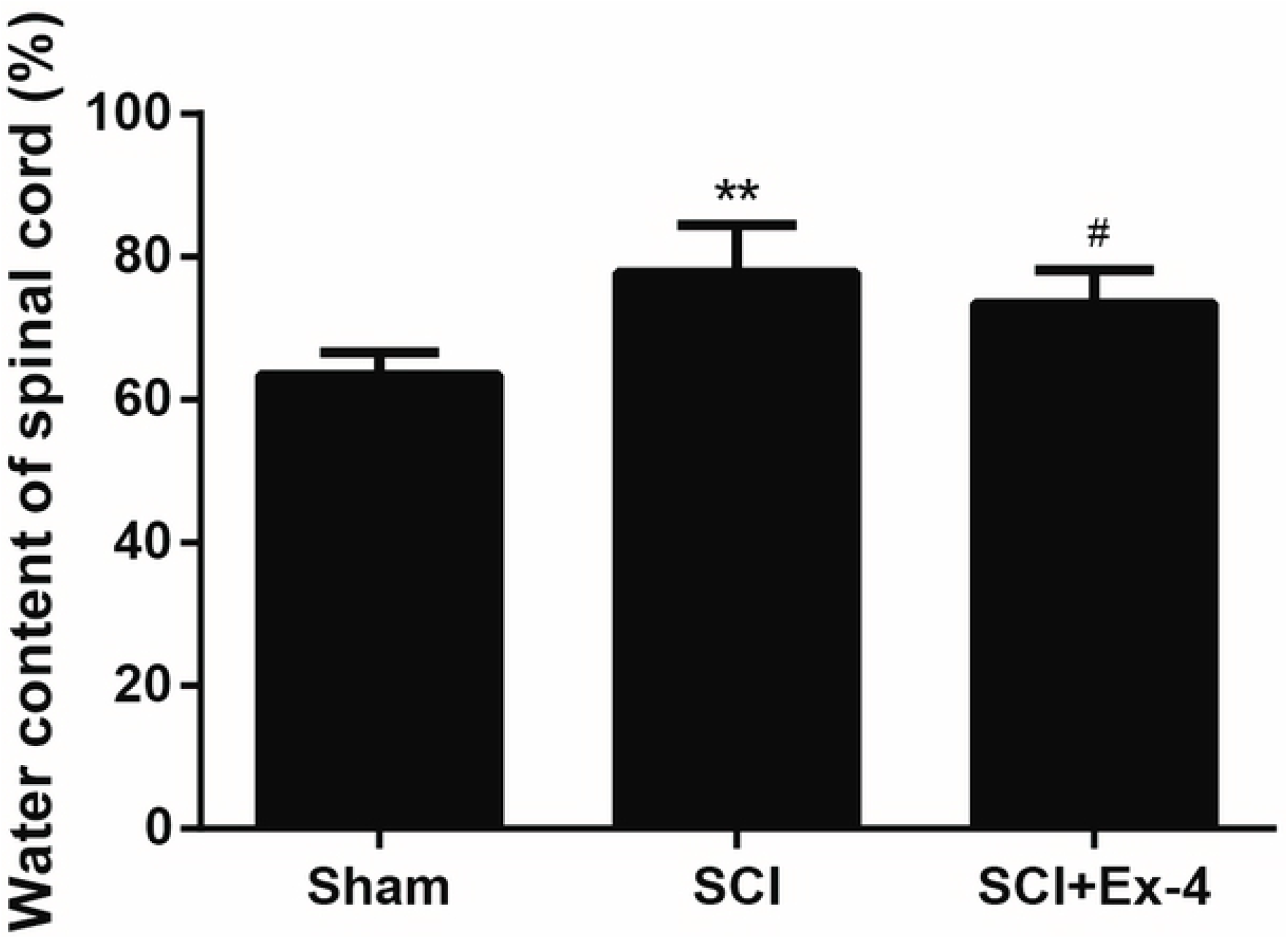
Water content of spinal cord in the three groups of rats, including a sham group, a SCI group and a SCI+Ex-4 group (n=12 per group) by wet/dry weight method. Ex-4treatment obviously alleviated edema of spinal cord in SCI rats. \*\**P*<0.01 vs sham group, ^#^*P*<0.05 vs SCI group.

### Exendin-4 suppresses inflammation of the injured spinal cord after SCI

Post-traumatic inflammatory reaction is a critical event in secondary damage after SCI (28). As shown in **Figure 3**, concentration level of IL-1α in SCI group was significantly increased than that in sham group (*P*<0.001). Moreover, compared with that in SCI group, concentration level of IL-1α in SCI+Ex-4 group was remarkably reduced after Ex-4 treatment (*P*<0.01). For the other cytokines (IL-1β, IL-6 and TNF-α), the results of above comparisons were similar as those of IL-1α.

**Figure 3.**
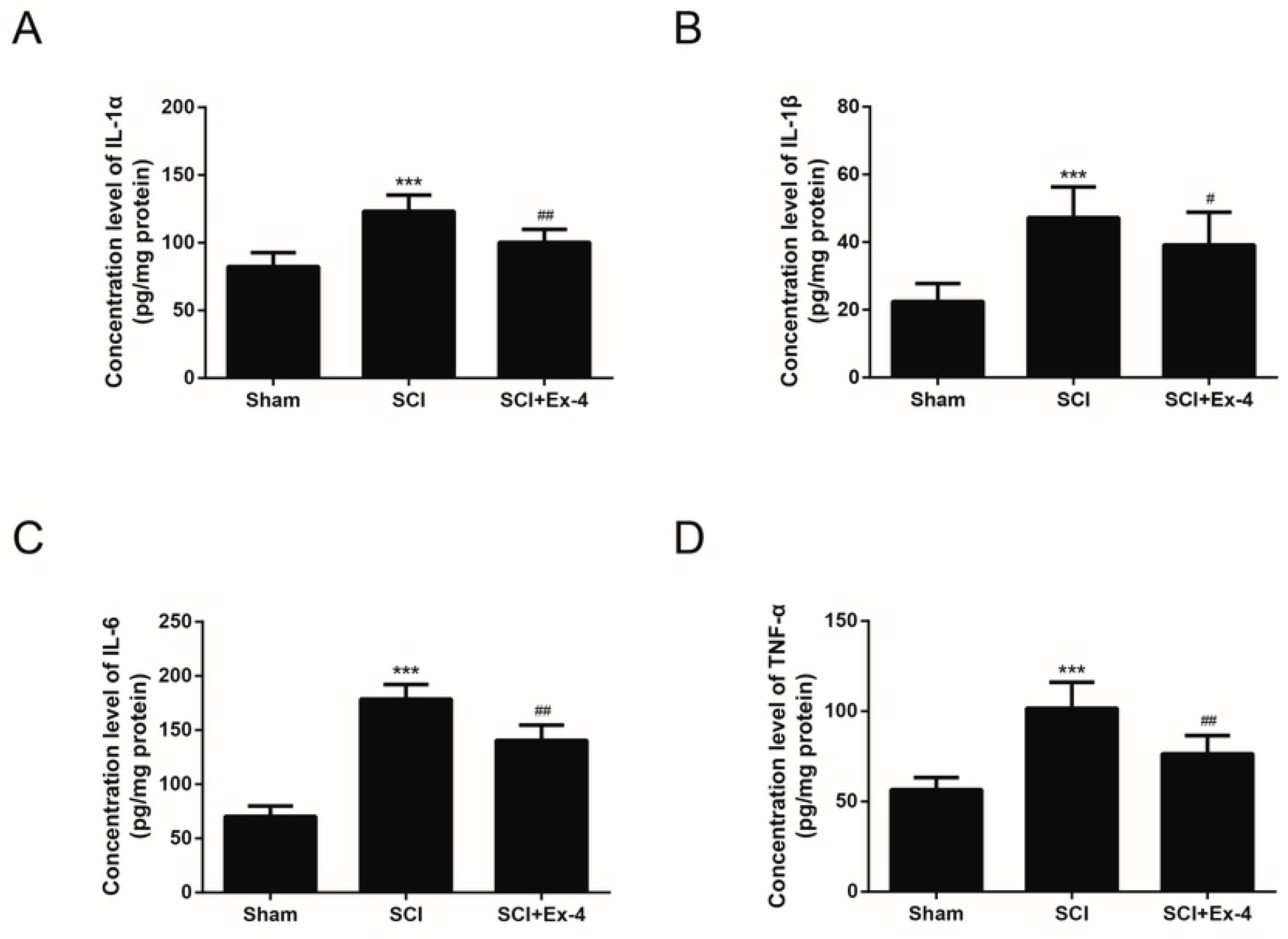
Concentrations of inflammatory cytokines in the three groups of rats, including a sham group, a SCI group and a SCI+Ex-4 group (n=12 per group). (A) Concentration levels of IL-1α; (B) Concentration levels of IL-1β; (C) Concentration levels of IL-6; (D) Concentration levels of TNF-α. \*\*\**P*<0.001 vs sham group; ^##^*P*<0.01 vs SCI group; ^#^*P*<0.05 vs SCI group.

### Exendin-4 inhibits cell apoptosis in the injured spinal cord after SCI

Apoptosis is a very important mechanism of secondary injury after SCI. As demonstrated in **Figure 4**, the cell apoptosis rate in the injured spinal cord of SCI group was significantly increased than that of sham group (*P*<0.001). Consistently, after Ex-4 administration, the cell apoptosis rate was remarkably decreased in comparison with that in SCI group, and the difference also had statistically significant (*P*<0.01). These results demonstrated that Ex-4 treatment could alleviate neurological deficits via inhibiting cell apoptosis in the injured spinal cord.

**Figure 4.**
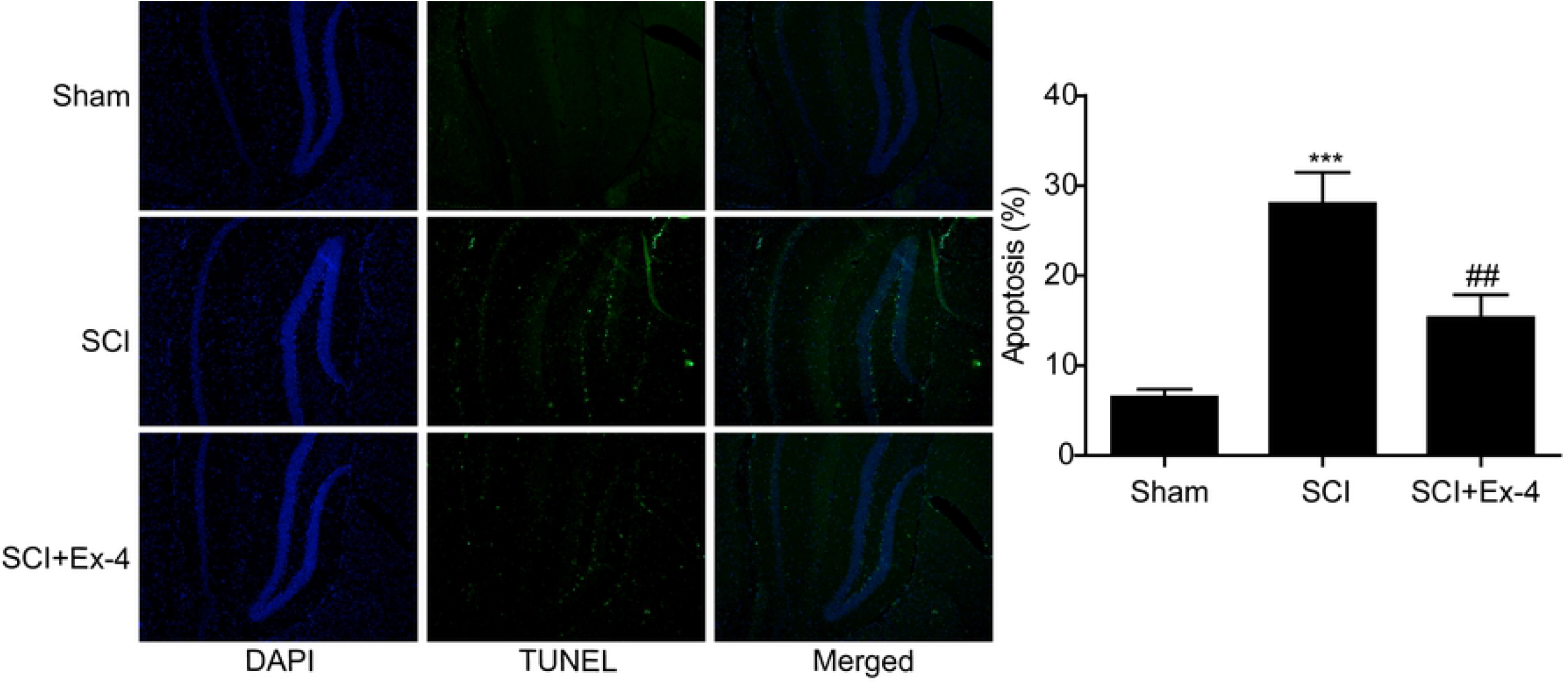
Cell apoptosis rates were detected through TUNEL assay in the three groups of rats, including a sham group, a SCI group and a SCI+Ex-4 group (n=12 per group). Ex-4 treatment obviously decreased the number of apoptotic cells after SCI. \*\*\**P*<0.001 vs sham group; ^###^*P*<0.001 vs SCI group.

### Exendin-4 inhibits PCBP2 and promotes Nrf-2 and OH-1 expression at mRNA and protein levels in the injured spinal cord after SCI

Actually, it still remains elusive how Ex-4 protects neurological function and inhibit cell apoptosis. In the present study, RT-qPCR analysis was performed to investigate whether mRNA expression of PCBP2, Nrf-2 and OH-1 had an association with SCI. As shown in **Figure 5A and 6A**, compared with that in sham group, the mRNA expression level of PCBP2 was significantly up-regulated and the levels of Nrf-2 and OH-1 were down-regulated after SCI (*P*<0.001). The mRNA expression level of PCBP2 was dramatically decreased in SCI+Ex-4 group than that in SCI group, and Nrf-2 and OH-1 were increased in SCI+EX-4 group (*P*<0.01).

**Figure 5.**
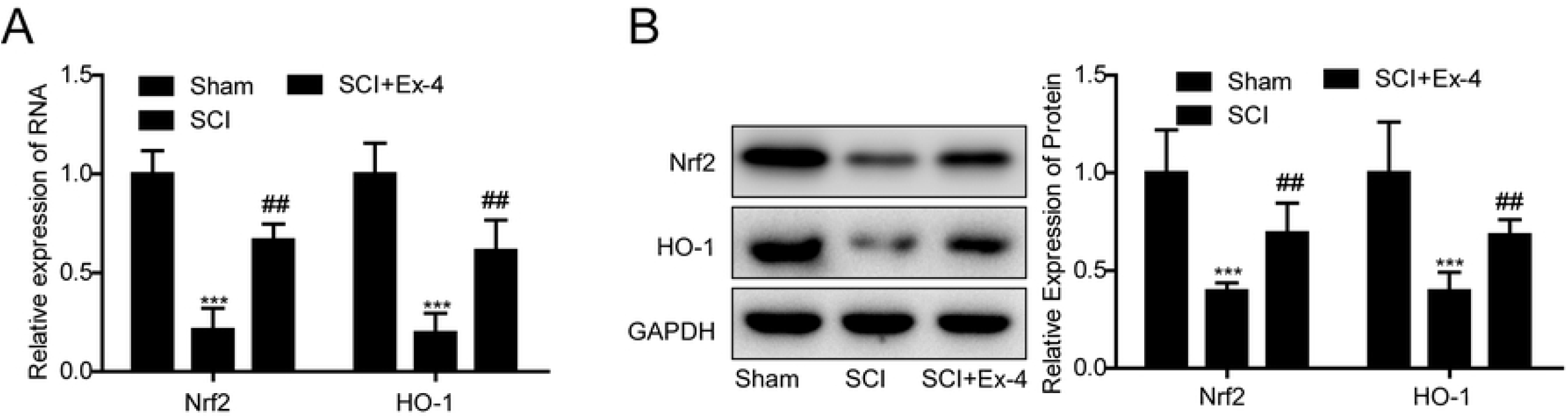
mRNA and protein levels of Nrf-2 / OH-1 were investigated in the three groups of rats, including a sham group, a SCI group and a SCI+Ex-4 group (n=12 per group). (A) mRNA level of Nrf-2 and OH-1 were significantly up-regulated after SCI, and they were down-regulated under the treatment of Ex-4. GAPDH mRNA was used as an internal control. (B) Protein level of Nrf-2 and OH-1 were significantly up-regulated after SCI and were down-regulated under the treatment of Ex-4. GAPDH protein was used as an internal control. \*\*\**P*<0.001 vs sham group; ^##^*P*<0.01 vs SCI group.

**Figure 6.**
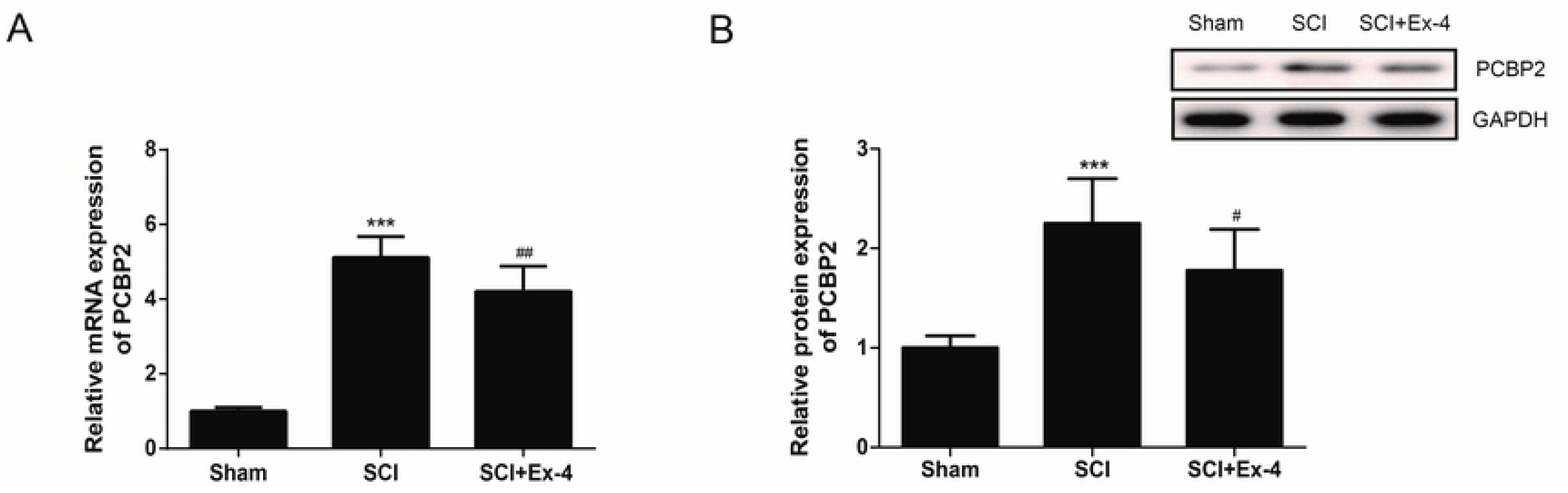
mRNA and protein levels of PCBP2 were investigated in the three groups of rats, including a sham group, a SCI group and a SCI+Ex-4 group (n=12 per group). (A) mRNA level of PCBP2 was significantly up-regulated after SCI and was down-regulated under the treatment of Ex-4. GAPDH mRNA was used as an internal control. (B) Protein level of PCBP2 was significantly up-regulated after SCI and was down-regulated under the treatment of Ex-4. GAPDH protein was used as an internal control. \*\*\**P*<0.001 vs sham group; ^##^*P*<0.01 vs SCI group.

Moreover, western blot assay was performed to detect the protein expression of PCBP2, Nrf-2 and OH-1 in SCI. As shown in **Figure 5B and 6B**, compared with that in sham group, the protein expression level of PCBP2 was significantly up-regulated and Nrf-2 and OH-1 were down-regulated after SCI (*P*<0.001). The protein expression level of PCBP2 was obviously decreased and Nrf-2 and OH-1 were increased in SCI+Ex-4 group than that in SCI group (*P*<0.01). There results indicated that Ex-4 might serve a neuro-protective role in SCI through inhibiting the expression of PCBP2 and promoting Nrf-2 / OH-1 pathway.

## Discussion

As we all know, SCI is a severe and complicated medical condition leads to lifelong neurological disabilities in both motor and sensory systems. To our knowledge, it was the first time to reveal that Ex-4 plays a role in promoting neurological function recovery and inhibiting neuronal apoptosis through effecting PCBP2 expression in SCI rat models.

Neurological impairment in SCI is mainly caused by two different mechanisms, including primary injury and secondary damage. The primary injury refers to neurological impairment caused by trauma, including neurons necrosis and axotomy, and the secondary damage occurs with a series of changes in cell metabolism and gene expression after primary injury, including edema, inflammation, ischemia, which eventually leads to neuronal apoptosis (29, 30). Therefore, neuroprotection and neurorecovery still remain the major strategies for SCI treatment.

An ideal neuroprotectant for treating SCI should be limited-toxic, easily delivered, and provide protection at all stages of injury. Ex-4, as a glucagon-like peptide-1(GLP) receptor agonist, can effectively down-regulate blood glucose level, stimulate pancreatic β-cell regeneration, induce transcription of pro-insulin gene and promote maturation and secretion of insulin (31). Ex-4 can also stimulate the growth and proliferation of human β-cells, and suppress cytokine-induced apoptosis (32). Fanet al. (33) demonstrated that Ex-4 could inhibit the apoptosis of retinal cell, balance the ratios of Bcl-2/Bax and Bcl-xL/Bax and decrease retinal reactive gliosis in diabetic Goto-Kakizaki rats. In addition, it was reported that Ex-4 serves a cardioprotective role, which could inhibit apoptosis of cardiomyocytes (34). Consistently, Gupta et al. (35) found that Ex-4 could inhibit cell apoptosis and promote the initiation of lipolysis, which played a protective role in ischemic injury of fatty liver. Moreover, increasing reports have demonstrated that Ex-4 could cross the blood-brain barrier (BBB) and act as neurotrophic factors in spinal cord and brain tissues through intraperitoneal administration (15, 36). Wang et al. (37) indicated that the ratio of cortical neuron apoptosis under oxygen/glucose deprivation (OGD) was obviously reduced after the treatment of Ex-4. Li et al. (15) found that Ex-4 treatment could reduce apoptosis of neuronal cell and ameliorate degeneration of motor neuron in amyotrophic lateral sclerosis. Of note, there are several similar conclusions about the association between Ex-4 and SCI. Li et al. (38) demonstrated that Ex-4 had a protective role in SCI rat models via inhibiting neuronal apoptosis and promoting the function recovery of motor nerve. Our present results extended previous findings by showing that SCI-induced hindlimb locomotion deficits were obviously ameliorated after Ex-4 treatment, indicating that Ex-4 could promote the recovery of locomotion recovery. Spinal cord water content was remarkably reduced and cell apoptosis in the spinal cord tissues was obviously inhibited by Ex-4 administration in SCI rats. All the results corroborate the conclusions of previous relevant studies. Morever, Ex-4 intervention could significantly promote the Nrf-2 / OH-1 pathway, which suggested that Ex-4 may alleviate oxidative stress and inflammatory responses by upregulating the Nrf-2 / OH-1 pathway, thereby inhibiting neuronal apoptosis.

PCBP2 is involved in regulation of gene expression at multiple levels, including transcription, processing, stability and translational regulation via regulating the binding capacity of poly (C). Mao et al. (24) found that PCBP2 plays an important role in neuronal apoptosis and astrocyte proliferation and PCBP2 knockdown obviously suppresses neuronal apoptosis after SCI. In our study, the levels of PCBP2 mRNA and protein were significantly up-regulated after Ex-4 treatment in SCI rat models, indicating a close association between Ex-4 administration and PCBP2 expression.

Taken together, in the present research we provided evidence that Ex-4 plays a crucial role in promoting the recovery of neurological function, alleviating the degree of edema of spinal cord and inhibiting cell apoptosis in the injured spinal cord after SCI, and PCBP2 might be a potential functional target of Ex-4 in SCI treatment.

## Acknowledgements

Not applicable.

## Ethics approval

The animal experiment was carried out according to the National Institute of Health’s Guidelines for the Care and Use of Laboratory Animals and was given permission by Animal Care and Research Committee of Sichuan Provincial People’s Hospital

## Authors’ contributions

Huaichao Luo, Qingwei Wang and Lei Wang designed experiments, analyzed data and interpreted results. Huaichao Luo and Qingwei Wang data acquisition. Huaichao Luo, Qingwei Wang and Lei Wang wrote the manuscript and prepared the figures. Huaichao Luo and Qingwei Wang reviewed and edited the manuscript. Lei Wang coordinated and directed the project. All authors approved the final version of the manuscript.

